# Eicosapentaenoic acid induces an anti-inflammatory transcriptomic landscape in T cells implicating a pathway independent of triglyceride lowering in cardiovascular risk reduction

**DOI:** 10.1101/2024.03.15.585315

**Authors:** Nathalie A. Reilly, Koen F. Dekkers, Jeroen Molenaar, Sinthuja Arumugam, Thomas B. Kuipers, Yavuz Ariyurek, Marten A. Hoeksema, J. Wouter Jukema, Bastiaan T. Heijmans

## Abstract

A twice-daily dose of highly purified eicosapentaenoic acid (EPA) reduces the risk of atherosclerotic cardiovascular disease among patients with high triglycerides and either known cardiovascular disease or those at high risk for developing it. However, the process by which EPA exerts its beneficial effects remains poorly understood. Here, we show that EPA can induce an anti-inflammatory transcriptional profile in non-activated CD4^+^ T cells. We find that EPA-exposed CD4^+^ T cells downregulate immune response related genes, such as *HLA-DRA, CD69*, and *IL2RA*, while upregulating genes involved in oxidative stress prevention, such as *NQO1*. Furthermore, transcription footprint analysis based on ATAC-sequencing reveals downregulation of GATA3 and PU.1, key transcription factors in T_H_2 and T_H_9 differentiation, and upregulation of REV-ERB, an antagonist of T_H_17 differentiation. By in parallel examining T cell responses to oleic acid, a monounsaturated fatty acid, and palmitic acid, a saturated fatty acid, we find that both the intensity of the transcriptomic response and the involvement of anti-inflammatory pathways is highly specific for EPA. Thus, EPA can induce an anti-inflammatory transcriptomic landscape in CD4^+^ T cells, a process that may contribute to the unexpectedly strong beneficial effects of EPA on the risk of atherosclerotic cardiovascular disease in clinical trials.

## Introduction

The risk of atherosclerotic cardiovascular disease (ASCVD) persists despite therapies that effectively control blood cholesterol levels including statins and PCSK9 inhibitors^1-3^. This residual risk has been attributed, in part, to elevated triglyceride levels in blood^4^. Nevertheless, most triglyceride influencing therapies, such as fibrates or niacin, have little cardiovascular benefit^5-9^. However, there is one triglyceride lowering drug that was found to strongly reduce ASCVD risk, namely, icosapent ethyl (IPE), which in the body is metabolized to eicosapentaenoic acid (EPA), a polyunsaturated fatty acid. The REDUCE-IT trial showed that patients who received 4g of IPE administered as 2g twice daily was superior to placebo in reducing triglycerides, cardiovascular events, and cardiovascular death among patients with high triglycerides and either known cardiovascular disease or those at high risk for developing it, and who were already on statin therapy with relatively well-controlled low density lipoprotein (LDL) levels^10^. The results of the trial and its interpretation has been much debated in literature^11, 12^. In particular, it remains largely unknown how EPA exerts its beneficial effects, and only limited studies have been carried out in model membranes or by examining whole blood^13-15^.

Atherosclerosis is regarded as a lipid-driven immune disease^16^. As such, the majority of immune cells in the atherosclerotic plaque are T cells, of which half are CD4^+17, 18^. Furthermore, CD4^+^ T cells aggravate atherosclerosis in established mouse models^19, 20^. Therefore, the study of CD4^+^ T cells is a promising route to further understanding ASCVD and investigating how EPA can influence these cells can indicate a potential mechanism underlying the beneficial effects of EPA on atherosclerosis. Interestingly, EPA was suggested to have anti-inflammatory properties as indicated by a reduction in CD4^+^ T cell proliferation, decreased differentiation towards T helper 1 (T_H_1) and T helper 17 (T_H_17), and increased or no effect on differentiation towards T helper 2 (T_H_2) and T regulatory (T_reg_) cells^21^. However, these studies were largely carried out in mouse models, or investigated *in vitro* during T cell activation, under polarizing conditions, or by measuring general T cell markers^22-28^. Thus, the effects of EPA on T cells remain incompletely understood and, in particular, it is unknown whether EPA can affect human CD4^+^ T cells in a non-activated state, as they occur in the circulation and where the primary interaction with EPA takes place.

We aimed to further elucidate the effects of EPA on CD4^+^ T cells by performing transcriptomic analysis on non-activated exposed cells. Furthermore, we assessed the specificity of the effects of EPA by exposing cells to two other fatty acids of different saturation, oleic acid (OA), a monounsaturated fatty acid, and palmitic acid (PA), a saturated fatty acid. To do so, we performed RNA and ATAC-sequencing on non-activated CD4^+^ T cells exposed to EPA, OA, PA, or control after 48h exposure. We show that EPA leads to a marked downregulation of many anti-inflammatory genes in non-activated CD4^+^ T cells as compared to control. The pronounced and specific effects on the transcriptomics landscape contrasted with the relatively modest effects of OA and PA.

## Methods

### Peripheral blood CD4^+^ T cell isolation and culture conditions

CD4^+^ T cell isolation and fatty acid exposure model were based on our previously described *in vitro* model with minor changes^29^. To obtain non-activated CD4^+^ T cells, peripheral blood mononuclear cells (PBMCs) were isolated from buffy coats of anonymous blood bank donors (Sanquin, Amsterdam, The Netherlands) by Ficoll paque (Apotheek LUMC, 97902861) gradient centrifugation. Next, CD4^+^ T cells were purified from the PBMCs using lyophilized human anti-CD4^+^ magnetically labeled microbeads (Miltenyi, 130-097-048) scaling the manufacturer’s instructions to ⅓ of the recommended volumes. CD4^+^ T cell purity was assessed on an LSR-II instrument at the Leiden University Medical Center Flow Cytometry Core Facility (https://www.lumc.nl/research/facilities/fcf/) with the BD FACSDiva™ v9.0 software (BD Biosciences). Cells were stained with anti-CD3-PE (BD Biosciences, 345765), anti-CD4-APC (BD Biosciences, 345771), anti-CD8-FITC (BD Biosciences, 555634), and anti-CD14-PEcy7 (BD Biosciences, 560919) and resuspended in 1% paraformaldehyde (Apotheek LUMC, 120810-001) to fix the cells prior to acquisition. Purity was >98% for all donors.

Prior to fatty acid exposure, ∼1*10^8^ isolated cells were cultured overnight to allow the cells to return to a resting state after the stress of the isolation procedure. This was done in T75 flasks (Greiner Bio-One, 658-175) at a density of ∼2.5*10^6^ cells/mL in 5% fetal calf serum (FCS) (Bodinco BDC, 16941) DMEM (Dulbecco’s Modified Eagle’s Serum (Sigma, 05796), 1% Pen-Strep (Lonza, DE17-602E), 1% GlutaMAX-1 (100x) (Gibco, 35050-038)) medium supplemented with 50 IU/mL IL-2 (Peprotech, 200-02) and incubated at 37°C under 5% CO_2_. To keep the cells in a non-activated state, no additional stimulus was added. Any CD4^+^ T cells not used directly after the isolation were kept in DMEM supplemented with 30% FCS, 1% Pen-Strep, 1% GlutaMAX-1, and 20% Dimethyl Sulfoxide (DMSO) (WAK-Chemie Medical GmbH, WAK-DMSO-10) medium at a density of ∼25*10^6^ cells/mL, and stored in liquid nitrogen.

Next, non-activated CD4^+^ T cells were cultured with either EPA (Cayman, 90110), OA (Sigma, O1383), or PA (Cayman, 10006627) for 48 hours at 37°C under 5% CO_2_. To this end, CD4^+^ T cells from each donor were plated in a 24 wells plate (density of ∼3.5*10^6^ cells/well) in 2mL 5% FCS DMEM for each condition (Fig. 1a). Cells were cultured in medium containing FCS to ensure cell viability during culture and to be more comparable to physiological conditions of the circulation where other lipids are also present. To assess the additional EPA, OA, or PA stimulus to the non-activated CD4^+^ T cells due to FCS in the culture medium, an FCS sample was measured via the Shotgun Lipidomics Assistant (SLA) method^30^ to estimate the fraction of fatty acids in the sample. The sample was prepped as previously described^31^ but with two modifications, a starting volume of 25μL FCS and 600μL MTBE was added instead of 575μL during the first extraction. Free EPA was 0.02μg/mL and EPA as components of larger molecules including cholesterol esters and sphingolipids was 0.13 μg/mL. Free OA was 0.29μg/mL and OA as components of larger molecules including cholesterol esters and sphingolipids was 4.93μg/mL. Free PA was 0.23μg/mL and PA as components of larger molecules including cholesterol esters and sphingolipids was 3.45μg/mL.

**FIGURE 1.**
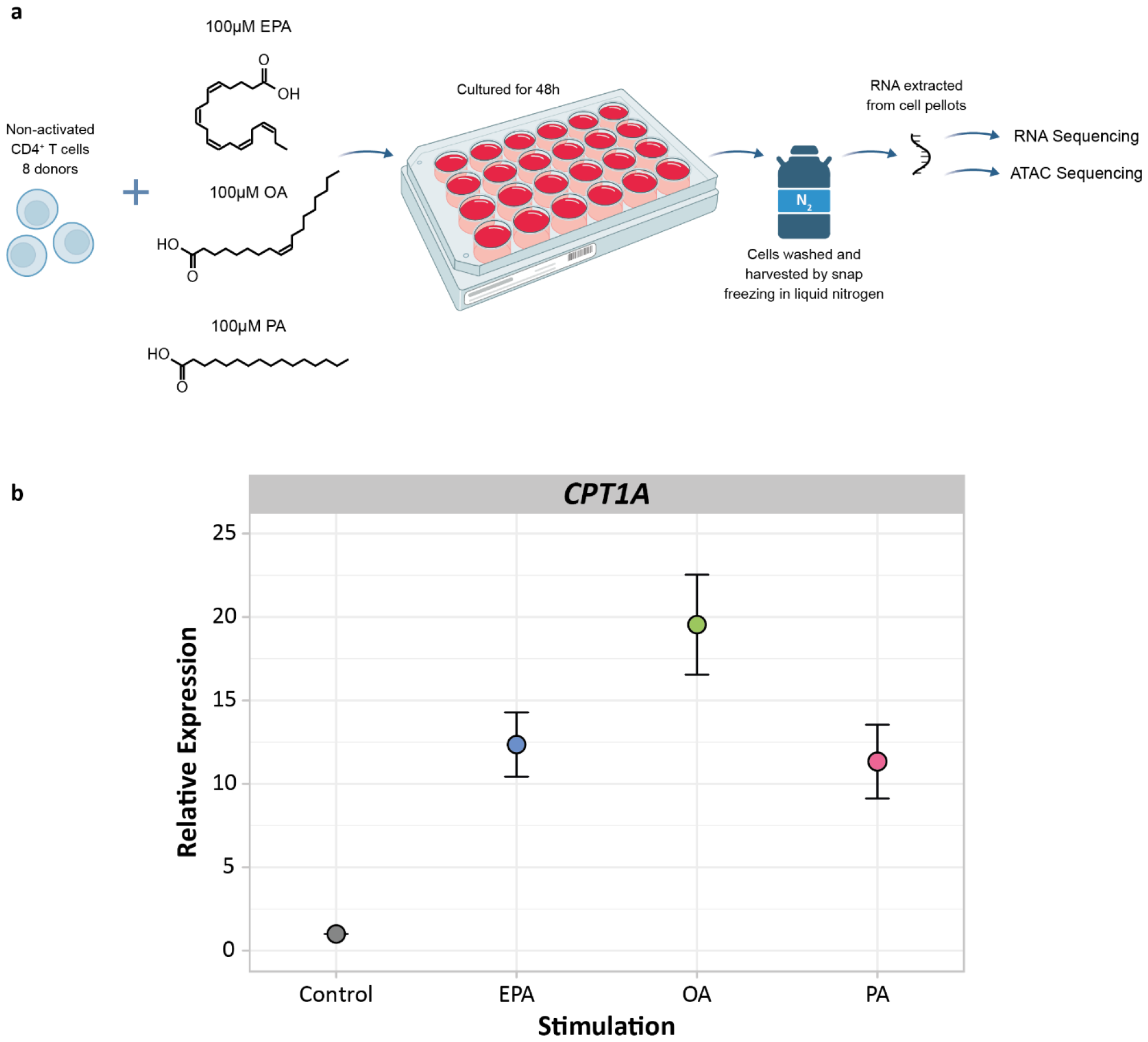
**(a)** Experimental set up for RNA and ATAC sequencing of EPA, OA, and PA exposed non-activated CD4^+^ T cells, n = 8. **(b)** Dot plot showing the relative expression of *CPT1A* after 48h of fatty acid exposure as a confirmation of the *in vitro* model by RT-qPCR. Values are colored by fatty acid. On average *CPT1A* was upregulated 12.4 SE 1.9 fold for EPA, 19.5 SE 3.0 fold for OA, and 11.3 SE 2.2 fold for PA, n = 8.

PA was dissolved in HPLC grade ethanol (Fisher Scientific, 64-17-5) to a final concentration of 5mg/mL to create a stock solution. The stock solution was vortexed briefly, sonicated in a sonicator (Branson, 2800) for 15 min, and heated for 15 min at 45°C. A small portion of the stock was extracted into a glass HPLC vial (Agilent Technologies, 5182-0714) to a final concentration of 5,000μg/mL. EPA and OA were dissolved from their stock in HPLC grade ethanol to a final concentration of 25,000 and 30,000μg/mL, respectively. The HPLC grade EtOH was then evaporated before the fatty acids were complexed to fatty acid-free (FAF) bovine serum albumin (BSA) (Sigma, A7030) in a 2% FAF BSA DMEM mixture (Dulbecco’s Modified Eagle’s Serum, 2% FAF BSA, 1% Pen-Strep, 1% GlutaMAX-1 (100x)) to a concentration of 151.25μg/mL for EPA, 141.25μg/mL for OA, and 128.2μg/mL for PA. Complexing fatty acids to BSA mimics physiological conditions as fatty acids are also bound to albumin in the human circulations^32^. Each fatty acid was further diluted to the final concentrations of 100μM (30.25μg/mL for EPA, 28.25μg/mL for OA, and 25.64μg/mL for PA) upon addition to the cells. The concentration tested was kept equal to ensure the cells were exposed the same amount of fatty acid particles and not influenced by concentration differences. Fatty acid stocks were stored under argon gas at −20°C to avoid oxidation.

As a control, HPLC grade EtOH was evaporated in a glass HPLC vial before adding 2% FAF BSA DMEM medium and added to the cells. The amount of 2% FAF BSA DMEM added to the wells was equal for each condition to keep the volumes equivalent. The CD4^+^ T cells were cultured for 48h at 37°C under 5% CO_2_. After exposure, the cells were washed and 1*10^5^ cells were used directly for ATAC sequencing preparation. Cells from the same donors for which ATAC sequencing was performed were later thawed from liquid nitrogen and exposed to the fatty acids as described previously. After 48h exposure, the cells were washed and 3*10^6^ cells were flash frozen in liquid nitrogen and stored at −80°C for RNA isolations. Cell viability and diameter were measured by Via1-Cassette™ (Chemometec, 941-0012) on a NucleoCounter® NC-200™ (Chemometec, 900-0200) and found to be > 95% and on average 9μm for each condition.

### RNA Isolation

To isolate total RNA for RNA sequencing and RT-qPCR, RNA was extracted from the cell samples using the Quick-RNA Microprep Kit (Zymo, R1050) according to manufacturer’s instructions. The RNA was quantified using a Qubit® 2.0 Fluorometer (Q32866) with the Qubit® RNA BR Assay Kit (Thermofisher, Q10211) according to manufacturer’s instructions. The RNA was placed over a second Zymo-Spin IC Column, washed, and a second DNase treatment performed to remove any residual DNA contamination from the samples. RNA integrity (RIN) values of the samples were on average 7.8 SE 0.1 as determined using an Agilent 2100 Bioanalyzer Instrument (G2939BA) with the Agilent RNA 6000 Nano Reagents (Agilent, 5067-1511). RNA was divided into two samples and stored at −80°C, 1μg for RNA sequencing and the rest for cDNA synthesis and RT-qPCR measurements.

### Real time-quantitative PCR

To measure the expression of *CPT1A* in all the cell samples, cDNA was synthesized with 200ng of the stored RNA using the Transcriptor First Strand cDNA Synthesis Kit (Roche, 04897030001) according to the manufacturer’s instructions. Quantitative real time PCR’s for *CPT1A* (Thermofisher, Hs00912671_m1, 4331182) were performed using the TaqMan™ Fast Advanced Master Mix (Thermofisher, 4444557) with 10ng cDNA per reaction on a QuantStudio 6 Real-Time PCR system (Applied Biosystems). All RT-qPCR reactions were performed in triplicate and outliers were removed if the Ct value measured differed more than 0.5% from the mean. Relative gene expression levels (-ΔCt) were calculated using the average of Ct values of *RPL13A* (Thermofisher, Hs03043887_gH, 4448892) and *SDHA* (Thermofisher, Hs00188166_m1, 4453320) as internal controls^33^. The fold change was determined using the 2^-ΔΔCt^ method, using the control as the reference. All statistical analyses were performed in R. Data are expressed as mean of the relative fold change and standard error. The reported P values were determined by applying a paired two-tailed student’s T test. P values < 0.05 were considered to be statistically significant.

### RNA Sequencing Analysis

RNA sequencing (RNA-seq) was performed to determine the differences in the transcriptome of control versus fatty acid exposed non-activated CD4^+^ T cells across time. The RNA from each of the samples was sent for sequencing (Macrogen, Amsterdam, NL). RNA-sequencing libraries were prepared from 200ng RNA using the Illumina Truseq stranded mRNA library prep (Illumina, 20020594) with a poly A selection. Both whole-transcriptome amplification and sequencing library preparations were performed in two 96-well plates with 26 samples in one plate and 6 in another. Quality control steps were included to determine total RNA quality and quantity, the optimal number of PCR preamplification cycles, and fragment size selection. No samples were eliminated from further downstream steps. Barcoded libraries were divided across two plates with 26 samples in one and 6 in the other and sequenced separately. Barcoded libraries were sequenced to a read depth of 20 million reads using the Novaseq 6000 (Illumina) to generate 100 base pair paired-end reads.

FastQ files are analyzed using the RNAseq pipeline (v5.0.0) from BioWDL (https://zenodo.org/record/5109461), developed by SASC (LUMC). The pipeline performed preprocessing on the FastQ files (including quality control, quality trimming, and adapter clipping), read mapping, and expression quantification. *FastQC* (v0.11.9) is used to check raw reads and *Cutadapt* (v2.10) to perform adapter clipping. Reads are mapped to a reference genome (Ensembl v105) using *STAR aligner* (v2.7.5a), and with *HTSeq Count* (v0.12.4) the number of assigned reads to genes per sample is determined.

Based on Ensembl gene biotype annotation, we included only protein coding genes for further downstream analysis (19,991 genes in total). We used the Bioconductor package *DESeq2*^34^ (v1.40.2) to test whether EPA, OA, or PA had an effect on gene expression as compared to the control. *DESeq2* fits a generalized linear model (GLM) assuming the negative binomial distribution for the counts. The model expresses the logarithm of the average of the counts in terms of one or more predictors. In this case, we used three models that had one of the fatty acids, subject identifier, and batch as predictors each. By including the subject identifier and batch in the models, we account for the dependence between measurements within the same subject and between different batches of sequencing^34^. Lowly expressed genes, i.e. that did not have at least a count of 1 in half of the samples per fatty acid and control, were removed, resulting in 12,938 genes for EPA, 12,949 genes for OA, and 12,971 genes for PA. The Benjamini-Hochberg procedure was used to correct for multiple testing at a false discovery rate (FDR) of 5%.

Differentially expressed genes per fatty acid were divided into upregulated or downregulated based on the log2 fold change values. 10 human pathway databases (BioPlanet 2019, WikiPathways 2019 Human, KEGG 2019 Human, Elsevier Pathway Collection, BioCarta 2015, Reactome 2016, HumanCyc 2016, NCI-Nature 2016, Panther 2016 and MSigDB Hallmark 2020) were queried using gene symbols, with 904 of 1170 queried genes for EPA, 51 of 60 queried genes for OA, and 26 of 33 queried genes for PA, present in at least 1 database. The identified clusters were then mapped for pathway enrichment using *clusterProfiler*^35^ (v4.8.3) with the background set to the 12,938 expressed genes for EPA, 12,949 expressed genes for OA, and 12,971 expressed genes for PA as determined above. Multiple testing correction using the Benjamini-Hochberg method at 5% FDR was performed over the combined results from the 10 databases. Pathways that included highly similar gene sets were grouped (Jaccard index > 0.7) and only the most significantly enriched pathway per group was retained.

### ATAC Sequencing Analysis

Post-exposure, the 1*10^5^ cells were taken off for ATAC sequencing and placed into DNA LoBind 1.5mL tubes (Eppendorf, 2231000945). The cells were washed 3x in ice cold buffered natrium chloride (PBS; pH 7.4; Fresen, 15360679). The samples were then handed off to the Leiden Genome Technology Center for library generation. The ATAC-sequencing libraries were generated using the Omni-ATAC protocol^36^. Briefly, the nuclei were isolated by lysing the cells in ATAC-Resuspension Buffer (RSB) (0.1% NP40 (Thermofisher, 85124), 0.1% Tween-20 (Thermofisher, 28320), and 0.01% digitonin (Promega, G9441)) for 3 min on ice. After washing the nuclei with 1mL wash buffer (RSB and 0.1% Tween) the nuclei were centrifuged for 10min at 4°C. After removing the supernatant, carefully avoiding the pelleted nuclei, the nuclei were resuspended in PBS. The nuclei were counted and normalized to 25,000 cells using the TC20 cell counter (BioRad, 1450102). The nuclei were combined with 25μL 2x TD buffer (TrisHCl pH 7.5 (Thermofisher, 15567027), NaCl (Thermofisher, A57006) and MgCl2 (Thermofisher, AM9530G)), 2μL Tn5 enzyme (Tn5 enzyme (Illumina, 15027865) and TD Tagment DNA Buffer (Illumina, 15027866)), 0.5μL 1% digitonin, 0.5μL 10% Tween-20 up to a volume of 50μL. The reaction was incubated at 37°C for 30min and then purified using AMPure Beads (Beckman Coulter, A63881) with a ratio of 1.8x and eluted in 10μL of EB (10mM Tris-HCl). The PCR was done using 2x Kapa HiFi Master mix (Roche, 09420398001) with the barcoded primers described in the Omni-ATAC protocol. After the PCR, the products were dual size selected using AMpure beads, first using 0.4x, followed directly by 1.2x. The ATAC-sequencing libraries were checked on the Femto Pulse (Agilent, M5330AA) and pooled equimolar for sequencing. No samples were eliminated from further downstream steps.

Barcoded libraries were sent for sequencing (Macrogen, Amsterdam, NL). An additional round of quality control was performed and the samples were then pooled and divided across one lane. Barcoded libraries were sequenced to a read depth of 30 million 150 base pair paired-end reads using the Novaseq 6000 (Illumina).

FastQ files were analyzed using the ChIP-seq pipeline from BioWDL (https://github.com/biowdl/ChIP-seq), developed by SASC (LUMC). The pipeline performed preprocessing on the FastQ files (including quality control, quality trimming, and adapter clipping), read mapping, and peak calling. *FastQC* (v0.11.9) is used to check raw reads and *Cutadapt* (v2.10) to perform adapter clipping. Reads are mapped to a reference genome (Encode GRCh38) using *BWA aligner* (v0.7.17), and *MACS2* (v2.1.2) was used to perform the peakcalling. These peak files were then processed using R (v4.3.0). Using *DiffBind* (v3.10.0), reads in the BAM files were counted for each peak. Next, the read counts per peak for each sample were merged to create one table containing all peaks and read counts of all the samples combined. *De novo* motif analysis was then performed using HOMER^37^.

### Data Availability

The data supporting the findings of this study are available within the article and its Supplementary information files. All other data including the raw files are available at the Gene Expression Omnibus repository, accession GEO (main combined submission: GSE254749, RNA sequencing submission: GSE254695, and ATAC sequencing submission: GSE254468).

## Results

### Transcriptomic analysis of EPA exposed non-activated CD4^+^ T cells

Non-activated CD4^+^ T cells were exposed to 100μM EPA, OA, or PA for 48h (n = 8, Fig. 1a). Exposure did not affect cell viability or diameter (Supp. Fig. 1a and b). To confirm a response by the cells due to the fatty acid exposure, the expression of *CPT1A*, the rate limiting enzyme in β-fatty acid oxidation, was measured. *CPT1A* expression increased as compared to control (EPA: 12.4-fold, SE 1.9; OA: 19.5-fold, SE 3.0; PA: 11.3-fold, SE 2.2; Fig. 1b). This signifies a consistent response to EPA, OA, and PA exposure.

Next, we studied the transcriptomic response of CD4^+^ T cells to EPA, OA, and PA using RNA-seq. The transcriptional response was compared to the control condition for each fatty acid. The number of differentially expressed genes (DEGs) and effect sizes were markedly larger for EPA, than for OA and PA (Fig. 2a) and there was limited overlap between the DEGs of each fatty acid (Fig. 2b). EPA induced 1170 DEGs (P_FDR_ < 0.05), 723 of which were downregulated and 447 of which were upregulated (Supp. Table 1a and b). In contrast, OA induced 60 DEGs (P_FDR_ < 0.05; 13 downregulated and 47 upregulated; Supp. Table 1c and d). PA induced found 33 DEGs (P_FDR_ < 0.05; 15 downregulated and 18 upregulated; Supp. Table 1e and f). Despite the high specificity of the transcriptional response of each fatty acid, 4 genes were upregulated upon exposure of all three fatty acids. These genes were involved in β-fatty acid oxidation (*CPT1A, SLC25A20, ACADVL*, and *ACAA2*) in line with a generic cellular response to fatty acid exposure regardless of the fatty acid type (Supp. Table 1g).

**FIGURE 2.**
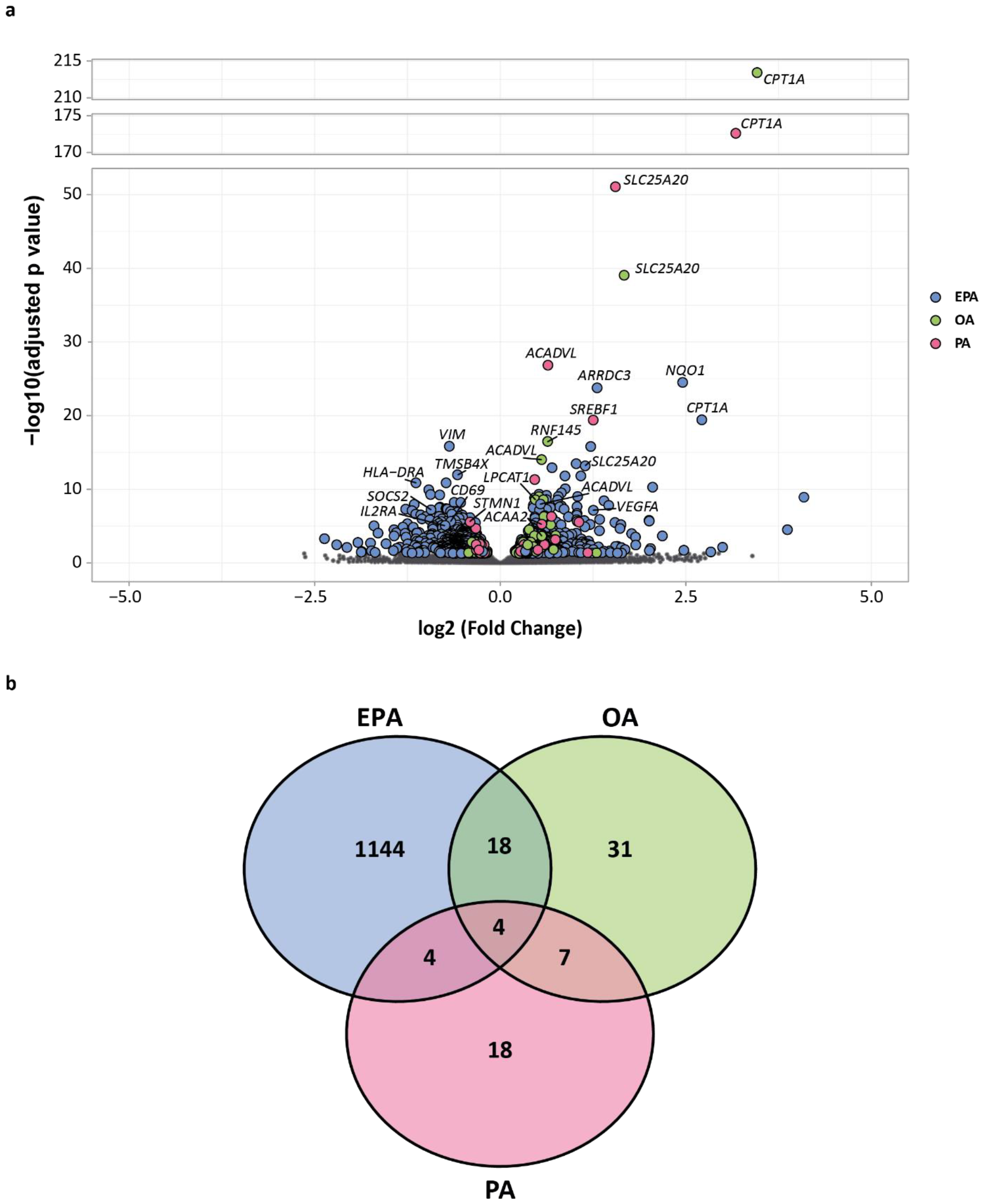
**(a)** Volcano plot showing the gene expression of non-activated CD4^+^ T cells exposed to either EPA, OA, or PA. All 19,991 protein coding genes are shown for each fatty acid. DEGs are colored by fatty acid and denoted by a larger size. Non-significant genes are shown in grey and denoted by a smaller size. Log2 fold change is used to show the direction of gene expression. **(b)** Venn diagram showing the unique response of non-activated CD4^+^ T cells to each fatty acid. Values are colored by fatty acid. There are 6 DEGs overlapping between all three fatty acids, 18 DEGs overlapping between EPA and OA, 4 DEGs overlapping between EPA and PA, and 7 DEGs overlapping between OA and PA.

We focused on the marked transcriptomic response of CD4^+^ T cells to EPA. Firstly, we analyzed the 723 downregulated genes in EPA exposed non-activated CD4^+^ T cells. The top three DEGs were *VIM* (vimentin), *TMSB4X* (thymosin beta 4 X-linked) and *HLA-DRA* (major histocompatibility complex, class II, DR alpha). *VIM* and *TMSB4X* both encode structures involved in the makeup of the cytoskeleton. HLA-DRA plays a central role in the immune response by presenting peptides to T cells. Remarkably, many other immune response genes were also downregulated, including *SOCS2* (suppressor of cytokine signaling 2), *CD69* (CD69 molecule), and *IL2RA* (interleukin 2 receptor subunit alpha). SOCS2 is a negative regulator of cytokine receptor signaling, particularly of IGF1R, an Insulin-Like Growth Factor whose expression is associated with the development of T_H_17 over T_reg_ subsets. CD69 plays an integral part in T cell activation, and IL2RA is an important regulator of T cell differentiation. A strong downregulation of immune-related processes was confirmed by a formal analysis of enriched biological processes. In particular, interleukin (IL)-2 signaling pathway (P_FDR_ < 0.001; 110 DEGs), antigen processing and presentation (P_FDR_ < 0.001; 27 DEGs), and interferon gamma response (P_FDR_ < 0.001; 47 DEGs) were enriched (Fig. 3a; Supp. Table 1h). This indicates that EPA reduces immune related gene expression in non-activated CD4^+^ T cells.

**FIGURE 3.**
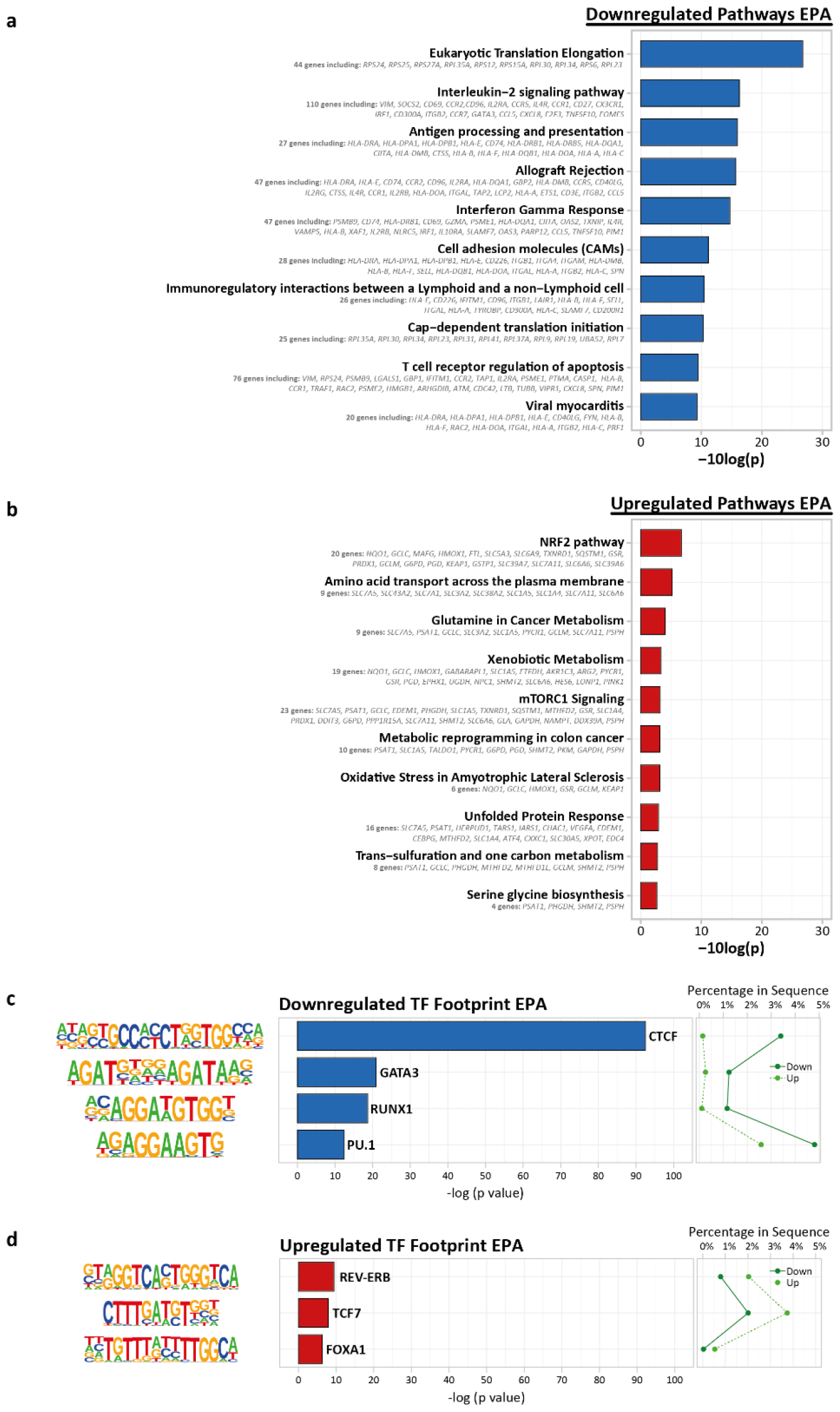
**(a)** Pathway enrichment analysis of all downregulated EPA DEGs generated using *clusterProfiler* using 10 human pathway databases. Top 10 enrichments are shown. **(b)** Pathway enrichment analysis of all upregulated EPA DEGs generated using *clusterProfiler* using 10 human pathway databases. Top 10 enrichments are shown. **(c)** Known motif analysis on promotors of down versus upregulated EPA ATAC peaks. Enrichment of transcription factor binding motifs was performed using HOMER. 4 motifs are shown with supplementing information on p-value, percentage of genes in upregulated gene set and percentage of genes in downregulated gene set, transcription factor name, -log(p-value), and percentage in sequence. **(d)** Known motif analysis on promotors of up versus downregulated EPA ATAC peaks. Enrichment of transcription factor binding motifs was performed using HOMER. 4 motifs are shown with supplementing information on p-value, percentage of genes in upregulated gene set and percentage of genes in downregulated gene set, transcription factor name, -log(p-value), and percentage in sequence.

Secondly, we analyzed the 447 upregulated genes in EPA exposed non-activated CD4^+^ T cells. The top three DEGs were *NQO1* (NAD(P)H quinone dehydrogenase 1), *ARRDC3* (arrestin domain containing 3) and *CPT1A*. NQO1 is involved in protecting cells against oxidative stress, which can be caused by lipid peroxidation and *ARRDC3* encodes a regulator of G protein-mediated signaling. Another gene of interest that was upregulated was *VEGFA* (vascular endothelial growth factor A). The enzyme encoded by this gene is a proangiogenic molecule known to be involved in creating immunosuppressive environments. This immunosuppressive profile was further supported by a formal analysis of enriched biological processes, which showed upregulation of the NRF2 pathway (P_FDR_ < 0.001; 20 DEGs; Fig. 3b; Supp. Table 1i). This pathway is the most important pathway for protecting cells against oxidative stress and has been shown to be involved in anti-inflammatory responses. Overall, these results suggests that EPA exposure can alter gene expression in non-activated T cells towards an anti-inflammatory profile by decreasing immune response related genes and pathways and increasing protective genes and pathways such as the NRF2 pathway.

Next, we investigated whether specific transcription factors may underlie the differential gene expression. To do this, we examined the enrichment of transcription factor binding motifs in loci that were more closed (down) versus more open (up) as determined by ATAC-sequencing. The top EPA downregulated motifs included CTCF, GATA3, RUNX1, and PU.1 (Fig. 3c; Supp. Table 1j). CTCF is a master regulator of chromatin looping and moreover, involved in effector cell differentiation^38, 39^. GATA3 and PU.1 are the key transcription factors for the development of T_H_2 and T_H_9 cells, respectively^40, 41^. RUNX1 is necessary for T cell maturation, knock outs of this transcription factor results in phenotypically and functionally immature T cells^42^. We next examined the enrichment of transcription factor binding motifs in upregulated versus downregulated genes. EPA upregulated motifs included, REV-ERB, TCF7, and FOXA1 (Fig. 3d; Supp. Table 1k). REV-ERB is an antagonist of RORγt, the key transcription factor for the development of T_H_17 cells^43^. TCF7 plays a role in the regulation of autoinflammatory T cell responses^44^. FOXA1 is involved in giving T_reg_ cells their suppressive properties^45^. These results further suggest that non-activated CD4^+^ T cells may decrease their ability to induce an immune response or effector T cell profile after EPA exposure.

### Transcriptomic analysis of OA and PA exposed non-activated CD4^+^ T cells

Non-activated CD4^+^ T cells were also exposed to either OA or PA and differential gene expression was measured. In line with our previous experiments, we found that OA exposure leads to downregulation of endogenous peptide antigen presentation (*HSPA5* and *PDIA3*; P_FDR_ < 0.001; 2 DEGs), electron transport chain and oxidative phosphorylation activity (*NDUFA12, NDUFB4*, and *ATP5F1C*; P_FDR_ < 0.001; 3 DEGs) and upregulation of cholesterol biosynthesis (*HMGCR, HMGCS1* and *DHCR24*; P_FDR_ < 0.001; 4 DEGs; Supp. Fig. 2a and b; Supp. Table 1l and m). Exposure to PA induced an opposite response, with the downregulation of cholesterol biosynthesis pathway (*HMGCR* and *SQLE*; P_FDR_ < 0.05; 2 DEGs), and upregulated beta fatty acid oxidation (*CPT1A, SLC25A20*, and *ACADVL*; P_FDR_ < 0.001; 3 DEGs; Supp. Fig. 2c and d; Supp. Table 1n and o). Thus, the changes in the transcriptome of OA and PA exposed cells seem to have a greater effect on cellular metabolism, particularly cholesterol metabolism, as compared to EPA.

The transcriptional responses observed were in line with the results of ATAC-sequencing based transcription factor footprint analysis. For OA only three motifs were downregulated including RAR:RXR, a motif known to play a part in the development of T_reg_ over T_H_17 cells (Supp. Fig. 3a; Supp. Table 1p). OA upregulated motifs included PU.1, as was found previously^29^ (accepted for publication in iScience) as well as IRF8, which is also involved in T_H_9 differentiation (Supp. Fig. 3b; Supp. Table 1q). PA downregulated motifs included IRF8 and GATA3 (Supp. Fig. 3c; Supp. Table 1r) and upregulated motifs included REV-ERB (Supp. Fig. 3d; Supp. Table 1s). OA and PA showed reversed effects on cholesterol metabolism processes which were mirrored in opposite associations with transcription factor binding motifs, indicating fatty-acid specific responses in non-activated CD4^+^ T cells.

## Discussion

IPE, the highly purified form of EPA, has been associated with reduced triglycerides, cardiovascular events, and cardiovascular death in individuals with relatively well controlled LDL levels, even when corrected for placebo response in the mineral oil control group, LDL, and CRP in the REDUCE-IT trial^10, 46-49^. The trials outcomes and interpretation have been widely debated and the mechanisms by which EPA exerts its beneficial effects remains incompletely understood^11, 12^. We show that EPA exposure can already produce distinct changes in T cells prior to activation by decreasing the expression of immune response genes and increasing the expression of genes involved in oxidative stress protection. This is further supported by changes in transcription factor binding sites in our ATAC-sequencing motif analysis, indicating a change in the epigenetic landscape of EPA exposed T cells. Furthermore, we show that EPA induces a unique response in non-activated CD4^+^ T cells as two other fatty acids of varying degrees of saturation, OA and PA, generated a smaller yet distinct effect on gene expression profiles in T cells as compared to control. Our findings imply that different fatty acids in the circulation can induce diverse effects on T cell transcriptomics, and that specifically EPA exposure may poise T cells to have clearer anti-inflammatory responses. These results underscore a potential mechanism by which EPA may mitigate ASCVD risk, suggesting its anti-inflammatory impact on T cells as a contributing factor. This is particularly noteworthy as T cells comprise over half of the immune cell population within atherosclerotic plaques^17, 18^.

Our results show that EPA exposure, but not OA nor PA, leads to a strong downregulation of immune response related genes and pathways. Particularly, antigen processing and presentation was downregulated in EPA exposed cells, denoted by, amongst others, the decreased expression of 14 different HLA genes. This gene group is crucial in inducing immune responses^50^. In addition, IL-2 signaling was also downregulated, which is required for T cell activation^51^. Downregulation of these pathways suggests the EPA exposed T cells may have a reduced ability to initiate an immune response, a key component of inflammatory responses in atherosclerotic plaques^52^. This result can support the finding that higher plasma EPA levels are associated with lower CVD risk in humans^53, 54^. Furthermore, pro-inflammatory pathways, such as interferon gamma response were downregulated in EPA exposed cells. IFNγ is primarily produced by pro-inflammatory T cell subset, T_H_1 cells, which have also been found to decrease upon EPA exposure^27, 55-57^. Moreover, the key transcription factors in T_H_2 and T_H_9 differentiation, GATA3 and PU.1, were also found to be decreased in our motif analysis. While T_H_2 cells have inconclusive effects on ASCVD, T_H_9 cells have been shown to aggravate it^58-60^. Thus, EPA exposure decreased immune response and pro-inflammatory pathways as well as suggests a reduced ability for key T cell differentiation transcription factors to bind.

In further support of EPA’s anti-inflammatory properties on non-activated CD4^+^ T cells, we found that the NRF2 pathway was upregulated upon EPA exposure. This pathway mainly functions in preventing oxidative stress in cells by activating genes involved in detoxification and removal of reactive oxygen species^61^. However, the NRF2 pathway has also been shown to aid in the anti-inflammatory responses of macrophages^62^ and has been suggested as a beneficial pleiotropic effect of statins^63^, as oxidative stress has been found to be a risk factor for ASCVD^64^. We also found an increased footprint for the transcription factors REV-ERB, TCF7, and FOXA1. These transcription factors are each involved in regulating T cell responses and generating a more anti-inflammatory T cell profile^43-45^. Overall, these data indicate that non-activated CD4^+^ T cells can already acquire an anti-inflammatory transcriptomic profile, which may play a role in the anti-inflammatory properties observed of EPA in clinical trials.

EPA has a distinct effect on CD4^+^ T cells. This is observed by our analysis of the effects of OA and PA on non-activated CD4^+^ T cells. The number of DEGs and effect sizes were smaller upon OA and PA exposure and distinctly different. Interestingly, OA and PA each had opposed effects on cholesterol biosynthesis, with OA upregulating and PA downregulating this pathway. Upregulation of cholesterol biosynthesis has been related to the development of T_H_17 cells by controlling RORγt activity, the key transcription factor in T_H_17 differentiation^65, 66^. This observation can be further supported by OA downregulating the RAR:RXR motif, which is involved in generating T_reg_ cells over T_H_17 and PA upregulating REV-ERB^43, 67^. The results of OA exposure also show the robustness of our approach as our findings here match what was found previously by our group^29^ (accepted for publication in iScience). These data suggest that our model is robust and each fatty acid induces its own unique response in non-activated CD4^+^ T cells.

We show that EPA exposure has beneficial anti-inflammatory effects on non-activated CD4^+^ T cells. This is relevant because T cells are largely non-activated in the circulation and it is in the circulation where T cells will encounter EPA when individuals are treated with IPE to reduce ASCVD risk. Nevertheless, our results are in line with experiments on activated T cells, which showed that EPA exposure decreased proliferation^22-26^, decreased T_H_1 and T_H_17 populations^27, 28^, and had no effect on or increased T_H_1 and T_reg_ populations^26-28^. Furthermore, our use of non-activated CD4^+^ T cells with no additional selection towards naïve, effector, memory, or specific T helper subsets, as well as in culture medium containing other lipids more closely represents the diversity of T cells and environment of the circulation in which EPA exposure takes place. Additionally, we utilized OA and PA, two fatty acids of varying degrees of saturation to establish the distinct effects of EPA. Nevertheless, this does not rule out that other fatty acids may have marked effects on non-activated CD4^+^ T cells as well^21^. A final limitation of our study is that we have employed an *in vitro* model and the effects of EPA on T cells *in vivo* should be studied in the context of trials of IPE. However, in mouse models, EPA supplementation has also been shown to reduce cholesterol levels^68^, whereas, in humans, the effects of EPA on ASCVD risk were independent of LDL lowering^47^. Therefore, using a validated *in vitro* model provides valuable insights to study the effects of EPA on human CD4^+^ T cells.

In conclusion, our data points to the fact that EPA produces a strong and specific anti-inflammatory transcriptional profile in non-activated CD4^+^ T cells comprised of both the downregulation of immune related genes and the upregulation of antioxidant genes. This profile is supported by transcription factor motif analysis and by the analysis of two other fatty acids of varying degrees of saturation. Our results contribute to the debate of how EPA exerts beneficial effects in human ASCVD. Our study gives an indication that the beneficial effects observed of EPA, as asserted in clinical trials, can already start in the circulation by inducing an anti-inflammatory transcriptional profile in non-activated T cells with potentially anti-atherosclerotic properties.

## Acknowledgements

The authors’ work is supported by the Dutch CardioVascular Alliance (The Dutch Heart Foundation, Dutch Federation of University Medical Centers, the Netherlands Organization for Health Research and Development, and the Royal Netherlands Academy of Sciences) for the GENIUSII project Generating the Best Evidence-Based Pharmaceutical Targets for Atherosclerosis (CVON2017-20).

## Author Contributions

B.T.H and J.W.J conceived the project. N.A.R. designed and conducted the experiments, analyzed the results, and drafted the manuscript. K.F.D. designed the analysis model and analyzed the RNA sequencing data. J.M. designed and performed *in vitro* model and prepped samples for ATAC sequencing. S. A. designed and performed *in vitro* model and prepped samples for RNA sequencing. T.K. aligned the RNA and ATAC sequencing data. Y.A. performed the ATAC sequencing library preparation. M.A.H. performed and analyzed the transcription factor footprint analysis. All authors contributed to the writing of the manuscript.

